# Balanced polymorphism in a floral transcription factor underlies an ancient rhythm of daily sex alternation in avocado

**DOI:** 10.64898/2025.12.22.695989

**Authors:** Jeffrey S. Groh, Marllon Fernando Soares dos Santos, Emmanuel Avila de Dios, Gracie Ackerman, Edwin Solares, Eric Focht, Danelle Seymour, Brandon S. Gaut, Mary Lu Arpaia, Graham Coop

**Affiliations:** Department of Evolution and Ecology, University of California, Davis, CA; Center for Population Biology, University of California, Davis, CA; Miller Institute for Basic Research in Science, University of California, Berkeley, CA; Botany and Plant Sciences Department, University of California, Riverside, CA; University of California, San Diego, CA; University of California, Irvine, CA

## Abstract

In avocado and certain wild relatives in Lauraceae, pollination occurs via a synchronized rhythm of floral sex timing between two hermaphroditic flowering types. A-type plants present female-phase flowers in the morning and male-phase flowers in the afternoon, while B-types show the complementary pattern, a form of heterodichogamy. We map this dimorphism in avocado to a genomic region overlapping a single strong candidate gene, SDMYB, where a dominant haplotype confers A-type flowering. SDMYB belongs to a subgroup of R2R3 MYB transcription factors established as key regulators of floral maturation in diverse species with links to circadian jasmonate signaling. Haplotypes at this locus form an ancient trans-species polymorphism maintained by negative frequency-dependent balancing selection over 44 million years, and they segregate in at least 26 non-avocado species, including in a genus where this mating system has not been reported. Across several species examined, rhythmic diel SDMYB expression is associated with biphasic floral anthesis, and the dominant allele, which contains nonsynonymous changes in conserved functional domains, exhibits a cis-regulated phase delay, corresponding to the delayed 2nd anthesis of A-types. The coupling of dichogamy with diel flower movements, widespread among magnoliids, is a likely precursor to daily forms of heterodichogamy. Absence of the SDMYB polymorphism in true cinnamon, which exhibits a highly similar mating system, suggests the possibility that heterodichogamy has convergently evolved within Lauraceae.

## Introduction

*“The daily rhythmic alternation of sexes… for the entire plant and the reciprocation of these changes in certain groups of plants reach a perfection of physiological regulation in avocados that is unapproached, as far as is now known, in any other group of plants.”*

— A.B. Stout, 1927

Flowers are a key innovation that have contributed to the rapid diversification and ecological dominance of angiosperms (Stebbins 1970; Endress 2011; Benton et al. 2022), and they play vital roles in agricultural systems and human cultures. Flowering time and maturation are tightly regulated by endogenous biological rhythms and coupled to seasonal and daily environmental rhythms to coordinate flowering with conditions that maximize reproductive success (Putterill et al. 2004; Creux and Harmer 2019). Considerable progress has been made in elaborating gene regulatory networks that regulate the transition to flowering, flower organ development, and flower maturation in model systems (Reeves et al. 2012; Bouché et al. 2016; Bowman and Moyroud 2024), yet connecting these findings to the genetic targets of mating adaptations in natural and agricultural systems remains an open challenge.

A widespread feature of angiosperm flowering is dichogamy - the temporal separation of male and female sexual maturity (Lloyd and Webb 1986; Bertin and Newman 1993). While dichogamy can limit pollen-pistil interference and promote cross-pollination, lineages in at least 14 angiosperm families have further evolved dimorphism for the direction of dichogamy (Renner 2001; Endress 2020; Garnock-Jones et al. 2025) . Such dimorphism, known as heterodichogamy, facilitates the evolution of a balanced sex ratio (Gleeson 1982), analogous to how Fisherian sex ratio selection maintains a 1:1 sex ratio in many species. Heterodichogamy polymorphisms are part of a broader class of mating-type polymorphisms where disassortative mating generates rare-mating-type advantage, a form of negative frequency-dependent balancing selection capable of maintaining genetic and phenotypic variation over deep evolutionary time scales. Well-studied examples like dioecy, heterostyly and self-incompatibility have contributed fundamental insights into the fitness advantages of outcrossing, the role of ecology in shaping mating strategies, the molecular basis of self-recognition, and the formation of supergenes (Darwin 1877; Charlesworth and Charlesworth 1979; Renner and Ricklefs 1995; Barrett 2010; Fujii et al. 2016; Gutíerrez-Valencia et al. 2021).

Heterodichogamy has evolved on both seasonal and daily timescales (Endress 2020). Recent work has begun to uncover the genetic basis of both seasonal and daily heterodichogamy, revealing a surprising diversity of genetic mechanisms that underlie its inheritance. Seasonal heterodichogamy is best known in Juglandaceae. At least four different genetic systems were recently shown to underlie the same phenotypic polymorphism present throughout most of the family (Groh et al. 2025a, Groh et al. 2025b, Liu et al. 2025), suggesting the possibility of turnover in the genetic mechanism. The functional changes involve genes linked to carbohydrate signaling and meristem regulation, and operate early in flower development, seemingly via small RNAs in at least two genera. The genetic basis of one form of daily heterodichogamy, flexistyly, was recently characterized in Zingiberaceae (Zhao et al. 2025). In this system, two morphs show synchronized patterns of style movement and anther dehiscence, controlled by rhythmic expression of a single gene with pleiotropic effects. Although seasonal and daily heterodichogamy operate on different developmental timescales, both are governed by Mendelian inheritance of oligogenic regions, suggesting repeatable genomic outcomes in the evolution of heterodichogamy from dichogamy. Aside from these few examples, however, the regulatory basis of heterodichogamy remains poorly known, and may reveal genetic constraints on sex ratio evolution in dichogamous populations.

The avocado (Persea americana) exhibits a remarkable example of heterodichogamy in its daily flowering rhythm (Fig. 1A). On a given tree, two sets of flowers will open (anthesis) and close synchronously at different times during a single day (Supplementary Videos 1-3). These two sets of flowers differ in their functional sex - one receiving pollen and the other releasing pollen. Thus, a hermaphroditic tree will be functionally male and female at different times of day. Avocado plants are further classified into two types, A and B, corresponding to whether the first set of flowers to undergo anthesis on a given day is functionally female (A-type) or male (B-type) (Stout 1927). Thus under typical conditions, A-type avocado varieties are functionally female in the morning and functionally male in the afternoon, while B-type varieties show the complementary pattern, promoting disassortative outcrossing between A- and B-type varieties.

**Figure 1:**
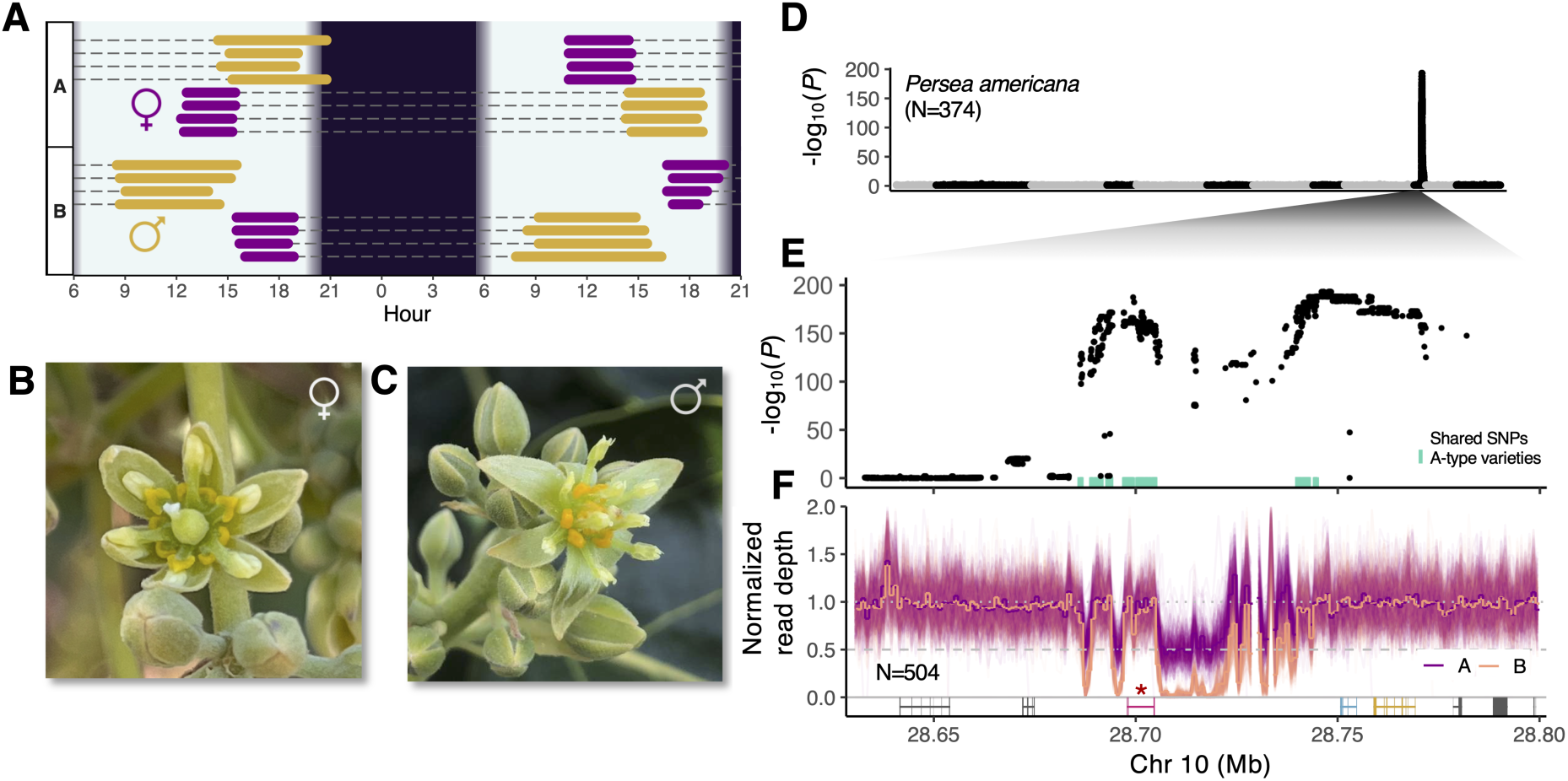
**A single-gene Mendelian locus controls synchronized dichogamy in avocado (*Persea americana* )**. **A)** Diagram of the synchronized dichogamy flowering pattern of avocado. Colored bars indicate the period when flowers are open in one of two sexual phases (purple - female, gold - male), and dotted gray lines represent when the same flowers are closed. Anther dehiscence occurs 1-2 hours after the onset of 2nd anthesis, so the period of pollen presentation begins later than indicated by the start of the gold bars. Top and bottom panels show data from representative flowers from a single individual each of A- and B-type, respectively, at UC Lindcove Research and Extension Center, Exeter, CA on April 10-11, 2024. **B,C)** Flowers of *P. americana* in female (B) and male phase (C). **D)** Association mapping for A- vs. B-type in a controlled cross. Alternating black and gray points represent SNPs on different chromosomes. **E)** Close-up view of the association peak. Colored tick marks at bottom indicate the locations of shared SNP polymorphisms among *Persea americana* A-type heterozygotes from an independent dataset. **F)** Normalized read depth in 1 kb windows for A- vs. B-types of avocado, with gene models shown at bottom and *SDMYB* marked by an asterisk. Photos in (B) and (C) by J.S.G. and M.F.S.S., respectively.

This alternating rhythm of sexual function at the level of individuals results from temporal offsets in a 2-day biphasic rhythm of individual flowers. Each individual flower is cosexual, opening first in the female phase, closing overnight, and opening the next day in the male phase. In A-type plants, the first anthesis occurs earlier on the first day, and the second anthesis occurs later on the second day, as compared to B-types (Fig. 1A). A large avocado tree may possess in excess of a million flowers, and the opening of flowers is staggered over successive days throughout the flowering season. This leads to a regular daily rhythm of alternating sexes in reversed directions between A- and B-types.

Since the original description of the avocado mating system (Stout 1927), highly similar mating systems have been reported in several species of Lauraceae, including other members of tribe Perseeae (Skutch 1945; Watanabe et al. 2016; Rogers 2024, Supplementary Videos 4-7), as well as in more distant relatives in other tribes including true cinnamon (Kubitzki and Kurz 1984; Hathurusinghe et al. 2023). These observations led some authors to suggest that this heterodichogamous mating system - termed synchronized dichogamy - might be ancestral to Lauraceae (Kubitzki and Kurz 1984), a large family originating in the Cretaceous whose ca. 3,000 modern representatives are almost almost all long-lived woody perennials distributed throughout the tropics and subtropics (Li et al. 2025). On the other hand, well-documented reports of synchronized dichogamy in Lauraceae remain sparse, and the occurrence of analogous types of heterodichogamy in other magnoliid families (Teichert et al. 2011; Endress and Lorence 2004; Endress 2020) suggests that this mating system might have instead repeatedly evolved within Lauraceae.

The avocado mating system has been both a point of fascination to botanists, as well as a challenge for growers, as both A- and B-type varieties are often planted together to maximize yield. The regularity and timing of the two phases is known to depend on both the diel light/dark cycle and temperature, contributing to variable crop yields of A- and B-types in different climates (Sedgley and Grant 1983; Sedgley 1985; Peterson 1956; Pattemore et al. 2018). Elucidation of the genetic regulation of the synchronized dichogamy in avocado could thus facilitate enhanced breeding efforts for a globally significant crop at the center of a multi-billion dollar industry. Previous work established that inheritance of dichogamy type in avocado is governed by a single locus with a dominant allele for A-type flowering (Ashworth et al. 2019). Here, we utilize a large mapping population of 374 resequenced avocado individuals together with de novo genome assemblies and time courses of gene expression from several species to identify the genomic and regulatory basis of synchronized dichogamy in avocado and its wild relatives. We find that synchronized dichogamy in avocado is linked to differences in rhythmic diel expression of alleles at a single flower-specific R2R3 MYB transcription factor, and that these alleles are both ancient and widespread throughout many species of Perseeae.

## Results

### Mapping the Synchronized Dichogamy locus in avocado

We performed a Genome Wide Association Study (GWAS) for A vs. B flowering type in 374 resequenced and imputed progeny from a controlled cross between the A-type heterozygote variety ‘Gem®’ (patent no. 3-29-5) and the B-type variety ‘Luna UCR®’ (patent no. BL516). This analysis revealed a single strong association peak on chromosome 10 (Fig. 1D-E) close to a known broad QTL peak for flowering type (Ashworth et al. 2019). For convenience, we refer to this association region as the SD-locus (for Synchronized Dichogamy), and to the dominant and recessive alleles as A1 and A2, respectively. In these and additional newly phenotyped and resequenced maternal half-sibs from the same parental varieties (N=504 total), A- and B- types show clear differences in average read depth against a haplotype-resolved assembly of Persea americana ‘Hass’ (Nath et al. 2022), an A-type heterozygote (Fig. 1F). These patterns indicated the presence of structural variation at the SD-locus and confirmed that B-type individuals from the controlled cross are recessive homozygotes (A2/A2 ) while A-type individuals are heterozygotes (A1/A2 ).

Three genes within the association peak were considered initial candidates (Fig. 1F, bottom in color). However, using published resequencing data from 23 additional avocado varieties (Solares et al. 2023; Rendón-Anaya et al. 2019), we found that the region of shared heterozygosity among a broader diversity of A-type varieties narrows the boundary of the SD-locus to a region overlapping only one of these genes (Fig. 1E, bottom). We identified this gene as a member of the flower-specific SG19 subfamily of R2R3 MYB transcription factors (Fig. S2A), which have been extensively characterized in model systems and shown to act as key regulators of floral opening, floral organ maturation, and the production of nectar, pigments, and volatiles (e.g. Sablowski et al. 1994; Liu et al. 2009; Colquhoun et al. 2011; Song et al. 2011; Reeves et al. 2012; Qi et al. 2015; Medina-Puche et al. 2015; Schubert et al. 2019; Chopy et al. 2023, and references within). Notably, this candidate gene in avocado is the closest homolog of the OsMYB8 gene in rice, where natural variation in the promoter region controls a difference in diurnal floret opening time between indica and japonica ecotypes (Gou et al. 2024). The avocado gene shows flower-specific expression, and is highly expressed in flowers (99.86*^th^* percentile among genes expressed in flowers), in clear contrast to the other two genes at the association peak (Fig. S2E,F). As variation at this gene is the strongest candidate for regulating synchronized dichogamy in avocado, we refer to it as SYNCHRONIZED DICHOGAMY MYB (SDMYB).

While the molecular function of SG19 R2R3 MYB transcription factors has not been previously investigated in any magnoliid to our knowledge, studies in eudicot and monocot systems have demonstrated that they localize to the nucleus and function as transcription factors (e.g. Spitzer-Rimon et al. 2010; Medina-Puche et al. 2015; Gou et al. 2024; Wu et al. 2021). Both the R2R3 DNA binding domain and the Transcriptional Activation Domain (TAD) are required for transcriptional activation of target genes (reviewed in Chopy et al. 2023). Consistent with this putative function, both avocado SDMYB alleles contain conserved R2R3 DNA binding domains and characteristic SG19 Transcriptional Activation Domains with a predicted alpha-helix (Liu et al. 2009) (Fig. S2A). The two avocado protein alleles have highly similar predicted 3D structures (Fig. S2B), yet notably differ by a single amino acid substitution at a strongly conserved site within the R2R3 DNA binding domain, and a frame-shift mutation within the TAD, suggesting possible functional changes. Given the distinct temporal flowering rhythms of A- and B-types, we hypothesized that these two SDMYB alleles should show distinct diel expression patterns.

### Temporal regulation of *SDMYB* underlies phase synchrony of A and B flowering types

To investigate the regulatory basis of synchronized dichogamy in avocado we constructed a time course of gene expression across the 2-day period of floral maturation beginning at the onset of first anthesis. To do this, we sampled flowers across developmental stages from 3 biological replicates of each morph every 3 hours spanning a 24 hr interval, after individually tagging flowers that had first opened the previous day (Supplementary Tables S1, S2). Using genome-wide transcript counts, we find all individuals show a clock-like oscillation along the first two principal components according to the time of day of sampling (Fig. 2A). Principal Component 1 (PC1, 17% of variance) captures expression variation associated with before noon vs. after noon while PC2 (15% of variance) captures variation associated with night vs. day. Thus, despite the conspicuous offset timing of sex expression between A- and B-types, the major axes of genome-wide expression variance in flowers reveal a largely shared diel trajectory. Consistent with this, we find that 47% of all flower-expressed genes (24% of all annotated nuclear genes) showed significant rhythmicity in both A- and B-types. Subsetting the data to only open flowers, PC2 separates samples in male vs. female phase (Fig. 2B), suggesting that the core transcriptional program regulating within-flower dichogamy is largely shared between types.

**Figure 2:**
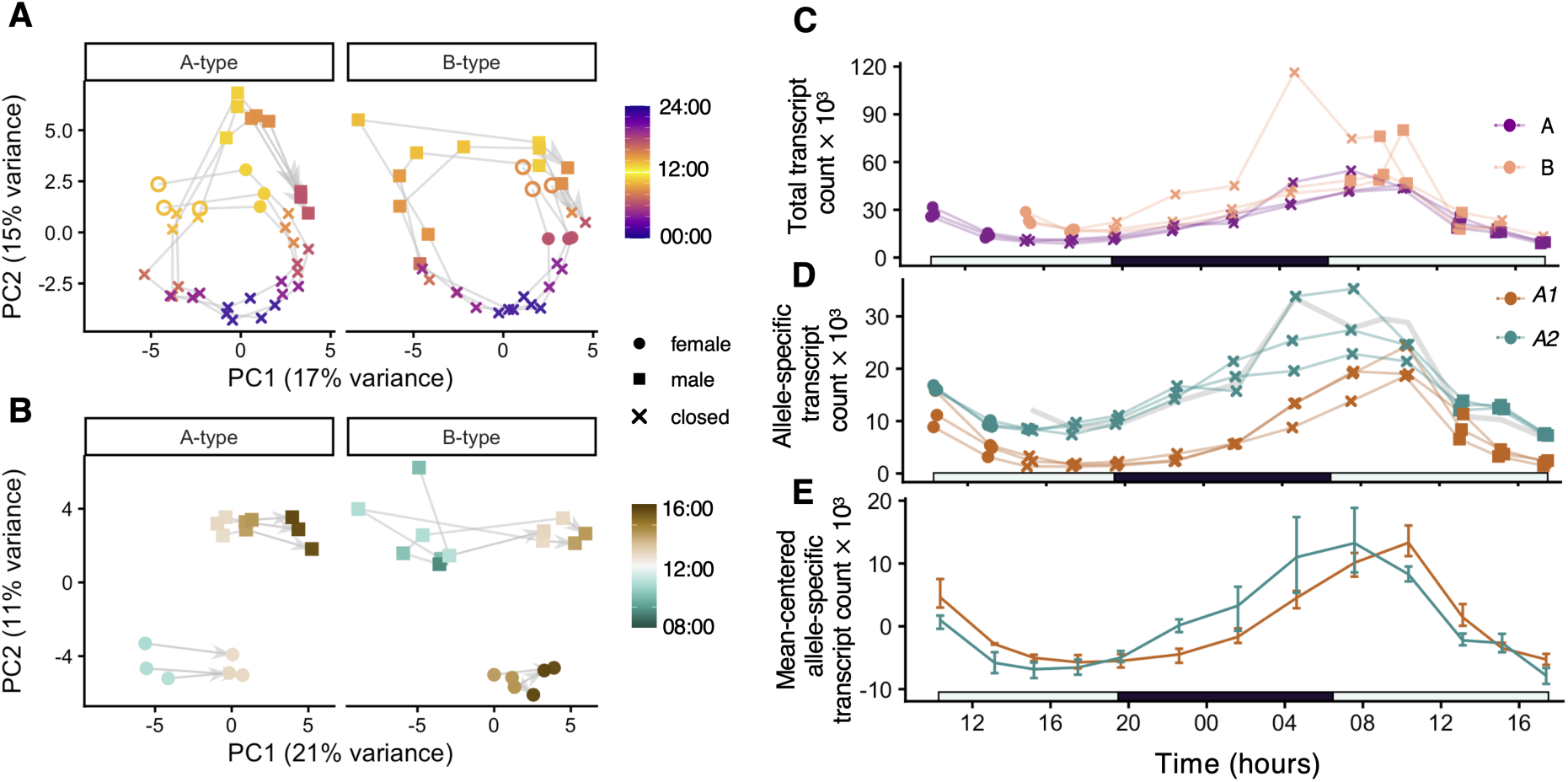
Temporal rhythms of avocado gene expression in flowers and allele-specific regulation at *SDMYB*. **A)** Principal Components Analysis (PCA) of genome-wide transcript counts. Each point represents an individual flower colored by the time of day of sampling, shape indicates developmental phase. Lines connect samples from the same individual, progressing along the developmental timecourse of flower maturation from open circle to arrowhead. **B)** PCA of transcript counts for only open flowers used in (A). **C)** Total expression of *SDMYB* transcripts following onset of first anthesis. Lines connect flowers from the same replicate. **D)** Allele-specific expression of *SDMYB* alleles in heterozygote A-type individuals of avocado, the same individuals shown in purple in (C) with each individual having two lines to represent both alleles. Gray line shows half of the average expression of *SDMYB* in B-type individuals from (C). **E)** Average of mean-centered allele trajectories of heterozygotes from (D) reveals a phase delay of the *A1* allele.

We found that SDMYB is differentially expressed between types across the time course (P=0.039) and exhibits striking temporal variation in allele-specific expression (Fig. 2C,D), in contrast to the other genes near the SD-locus (Fig. S2F-G), further supporting our identification of SDMYB as the best functional candidate. In both avocado types, total SDMYB expression declines immediately following first anthesis, then beginning in the afternoon gradually increases towards a local maximum just prior to or coincident with second anthesis on the following day (Fig. 2C). Although we could not determine the expression level of SDMYB in flowers prior to their first anthesis (the first sampling point corresponds to the start of first anthesis), our data appear consistent with a model whereby SDMYB expression must reach a threshold level for the flower to open, mirroring a pattern seen for EOBII, a Petunia ortholog of SDMYB (Colquhoun et al. 2011).

Total SDMYB expression is on average lower in A-types compared to B-types (Fig. 2C, P= 0.035, χ^2^ = 4.28, d.f.=1). This difference is accounted for by lower expression of the A1 allele in heterozygotes (Fig. 2D, P< 2.2 ⇥ 10^16^, χ^2^ = 141.8, d.f.=1), whereas the A2 allele in heterozygotes shows expression levels largely consistent with half of that seen in A2/A2 B-types (Fig. 2D, gray line). These patterns imply there are cis-acting regulatory differences between the haplotypes. However, we noted a significant difference in expression between A1 in A-types and the haploid equivalent in B-types at the first time point (P=0.019, Bonferroni corrected), suggesting the regulation of A2 may be altered in a heterozygous background. In addition to the difference in average expression of the alleles, we detected a phase shift between the two alleles, with the A1 allele showing an average 2 hr phase delay in heterozygotes (Fig. 2E, cosinor regression P=1.4 ⇥ 10*^—^*^4^). As this expression delay closely corresponds to the delayed timing of 2*^nd^* anthesis of A-types, these data strongly suggest that avocado synchronized dichogamy is determined in part by the differential temporal regulation of SDMYB alleles controlled by variation in cis.

### The SD-locus is a trans-species balanced polymorphism

Synchronized dichogamy has been reported in several members of Lauraceae, including in close relatives of avocado in Persea subg. Eriodaphne and Machilus (tribe Perseeae) (Skutch 1945; Watanabe et al. 2016) and in more distant relatives in tribes Cinnamomeae and Mezilaureae (Kubitzki and Kurz 1984; Hathurusinghe et al. 2023). To identify whether the avocado SD-locus might also regulate this mating system in other species, we first generated haplotype-resolved genome assemblies (Table S3) from phenotyped individuals of M. thunbergii (A-type, Supplementary Video 4) and P. caerulea (A-type, Supplementary Video 6), as well as an individual of P. podadenia (unknown type), together with P. americana ‘Puebla’, another A-type avocado variety. We used these together with published Lauraceae genome assemblies and both published and new resequencing data (Table S4) to better resolve species relationships within Perseeae, and to provide a phylogenetic framework for probing the evolutionary history of the SD-locus polymorphism.

Phylogenetic analysis using 7,842 single copy orthologs confirmed reciprocal monophyly of the three accepted Lauraceae tribes represented in our dataset (Fig. 3A, S3). However, our phylogeny demonstrates unambiguously that Persea is not monophyletic, which has been previously suggested (Rojas et al. 2007; Rohwer et al. 2009). While a taxonomic revision of Persea is not our focus, our analysis appears consistent with the suggestion that Persea subg. Persea and Persea subg. Eriodaphne should be treated as separate monophyletic genera (Rojas et al. 2007).

**Figure 3:**
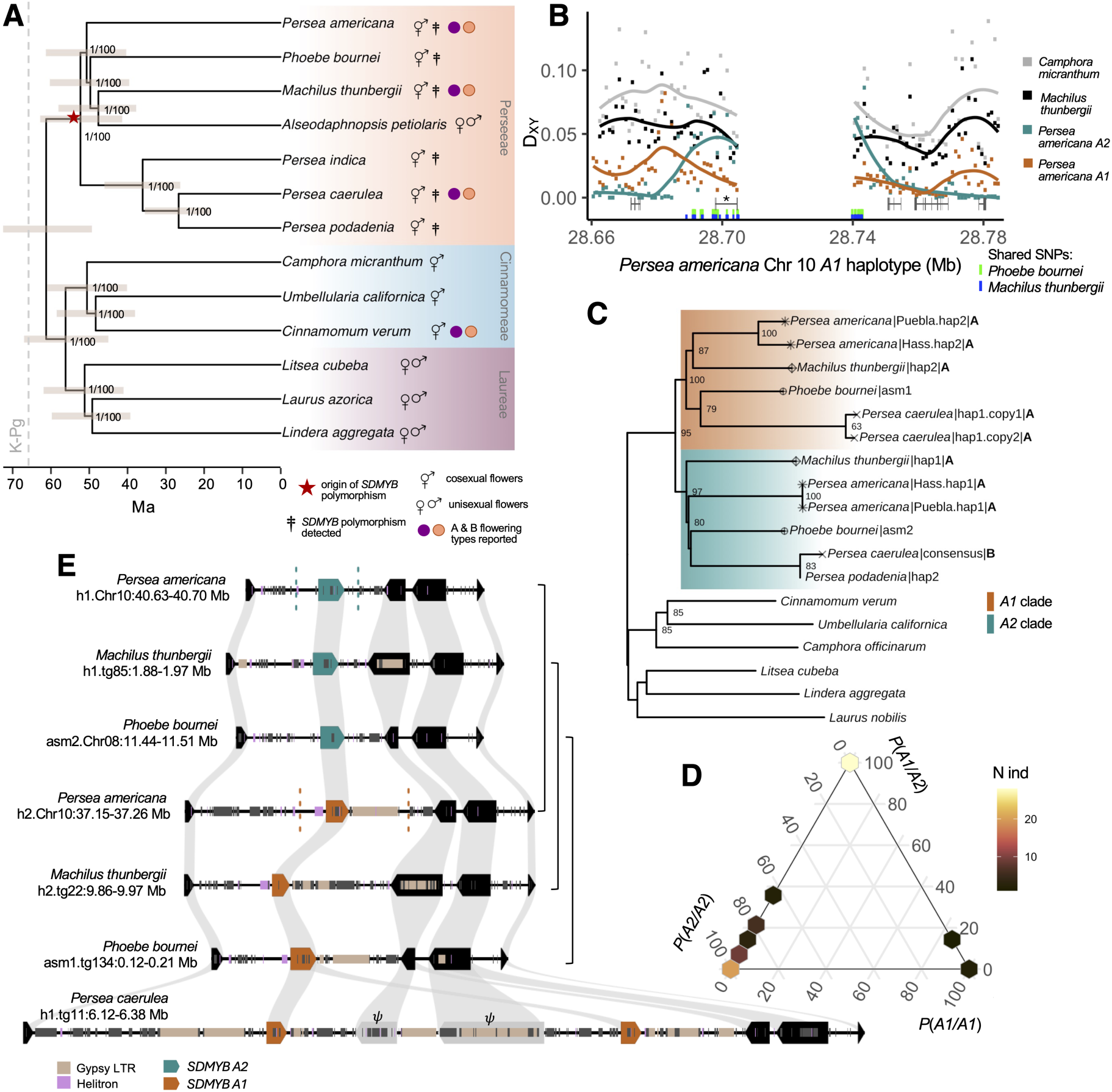
**Trans-species polymorphism and structural variation of SD-locus haplotypes**. **A)** A phylogeny of three tribes of Lauraceae (see also Fig. S3). Axis at bottom indicates timescale, and gray error bars represent 95% HPD intervals. Internal nodes are labeled with Local Posterior Probability support values / bootstrap support values. Red star indicates approximate divergence time of the *SDMYB* alleles. **B)** Nucleotide divergence between avocado (*A1* haplotype) and other Lauraceae genomes. Squares indicate values measured in 1 kb windows, and lines show LOESS smoothed curves. Gene models are shown at bottom (*SDMYB* denoted with asterisk) and the locations of shared trans-species SNPs between avocado and *Phoebe bournei* / *Machilus thunbergii* are indicated in colored tick marks. **C)** Maximum likelihood phylogeny of the SD-locus, rooted with *Sextonia rubra*. Node labels indicate bootstrap support, support values below 60% not shown. Tip shapes highlight species segregating for the trans-species polymorphism. Where known, flowering types (A/B) are indicated after species name and assembly identifier. **D)** Ternary plot showing genotype posterior probabilities for the SD-locus for 72 individuals representing 56 species of Perseeae. Hexagons indicate the locations of data points, with color indicating the number of individuals at a location. **E)** Synteny plot of the local region around *SDMYB* for assemblies from several Perseeae species. Pointed rectangles indicate gene models and direction of transcription, with ribbons connecting orthologs. *SDMYB* is colored by allele (orange - *A1*, blue - *A2* ). Narrow rectangles along haplotypes show repeat annotations, with two particular classes highlighted in color. Braces at right connect haplotypes from the same species. Dotted lines on *P. americana* haplotypes indicate the region of trans-species polymorphism surrounding *SDMYB*. in the *P. caerulea* assembly denotes predicted pseudogenes.

We first examined the SD-locus in close relatives of avocado known to be phenotypically polymorphic for A- and B-types - Machilus thunbergii (Watanabe et al. 2016), and Persea caerulea (Skutch 1945). Between the two assembled haplotypes of the A-type Machilus thunbergii, we observed a high concentration of shared heterozygous sites with A-type avocado varieties (trans-species polymorphisms) closely aligning with the boundaries of the avocado SD-locus (Fig. 3B, bottom, blue). We next determined that our A-type P. caerulea assembly individual was largely homozygous for A1 variants at these same sites, consistent with it having an A1/A1 genotype. RNA-seq data from a separate A-type individual of this species showed both A1 and A2 variants at these sites, while data from a B-type individual contained only A2 variants, confirming that the avocado SD-locus SNPs segregate in P. caerulea with the same phenotypic association. The nonsynonymous substitution within the the R2R3 DNA binding domain as well as the frame-shift mutation within the TAD of SDMYB are conserved differences between the A1 and A2 haplotypes across these species (Fig. S2A), suggesting these arose early in the formation of the SD-locus haplotypes.

The narrow region of phenotype-associated trans-species polymorphism suggested these haplotypes are maintained by long-term balancing selection. Nucleotide divergence between the avocado haplotypes sharply increases at the boundaries of the SD-locus, reaching a level comparable to the divergence of Persea americana and Machilus thunbergii (Fig. 3B), suggesting the haplotypes could have diverged in the common ancestor Perseeae. A maximum likelihood phylogeny of the SD-locus supported that the sets of A1 and A2 haplotypes from across Perseeae are reciprocally monophyletic sister groups (99% and 88% bootstrap support) (Fig. 3C). To calibrate the age of their divergence, we time calibrated our core Lauraceae phylogeny grafted into a magnoliid backbone tree (Fig. S3), leveraging information from several fossil calibrations (Tables S5-6). Doing so, we infer an Eocene origin of tribe Perseeae (52.2 Ma, 95% HPD [39.5, 60.2]). Using the divergence of Perseeae from the Cinnamomeae-Laureae clade to calibrate the divergence of the SD-locus haplotypes, we obtain an estimate of 44.1 Ma, 95% HPD [35.4, 52.2]. This estimate is consistent with these haplotypes originating in the common ancestor of Perseeae, or possibly introgression in the very early history of this group.

Long-term maintenance of the SD-locus haplotypes by balancing selection since the common ancestor of Perseeae predicts that they should segregate in many of the ca. 400 species of Perseeae distributed throughout the tropics and subtropics. To test this hypothesis, we inferred SD-locus genotypes for a set of 72 individuals representing 56 species in Perseeae using published and newly generated sequence data (Table S7), together with our identification of diagnostic SD-locus polymorphisms within SDMYB exons (Fig. S2). Doing so, we identified SD-locus haplotypes segregating in 26 non-avocado species (Fig. 3D, Table S7). The signal of shared exonic polymorphisms we detect is restricted to SDMYB and not seen in surrounding genes (Fig. S5). In this cross species sample, A1/A2 and A2/A2 genotypes are both common (53% and 43%, respectively) whereas A1/A1 homozygotes are rare (4%), consistent with the prediction that the SD-locus polymorphism is selectively maintained through disassortative mating in most of these species. Notably, we also identified A1 and A2 haplotypes contained within two independent genome assemblies from Phoebe bournei (Chen et al. 2020a; Han et al. 2022) (Fig. 3E), and confirmed that the individual from which the A1 haplotype assembly was heterozygous across most of the trans-species polymorphic SNPs identified at the SD-locus (Fig. 3B, bottom, green).

While synchronized dichogamy is well-documented in the Lauraceae tribe Cinnamomeae (Hathurusinghe et al. 2023)and has also been reported in tribe Mezilaureae (Kubitzki and Kurz 1984), our phylogeny inference (Fig. 3C) and divergence across the region (Fig. 3B) indicates that the SD-locus haplotypes are not as old as the divergence of Perseeae from other Lauraceae tribes. Further supporting this, we did not see the diagnostic SDMYB SNPs segregating in resequencing data from three individuals of true cinnamon (Chen et al. 2020b) (C. verum), where synchronized dichogamy is well-documented (Hathurusinghe et al. 2023), nor in resequencing data from 24 additional species in tribe Cinnamomeae (Wang et al. 2022). Finally, we examined a natural population of Umbelullaria californica (California bay laurel) and found that it does not exhibit synchronized dichogamy (Fig. S6). Altogether, these results indicate that the SD-locus polymorphism is not found within tribe Cinnamomeae and that synchronized dichogamy is not ubiquitous in this group, suggesting there could have been multiple independent origins of synchronized dichogamy within Lauraceae.

Haplotype-resolved genome assemblies revealed that structural differences between the A1 and A2 haplotypes are a shared feature across Perseeae species (Figs. 3E, S4A-C). Dominant A1 haplotypes show a moderate increase in length compared to A2, in part accounted for by transposable elements, most notably Gypsy-LTR-derived sequence downstream of SDMYB. In the A1/A1 P. caerulea homozygote, both haplotypes contain a tandem duplication of the A1 allele of SDMYB, which we did not detect in a RNAseq data from a B-type P. caerulea individual nor in the A2 long-read haplotypes from our P. podadenia assembly individual, a close relative. While these SDMYB paralogs show identical coding sequence, two other genes in the duplicated segment appear pseudogenized (Fig. S4E), suggesting that both SDMYB paralogs are selectively maintained. Despite the observed structural differences between haplotypes, it is noteworthy that the dichogamy-linked region has not expanded into a large heteromorphic region, mirroring an emerging trend identified in other heterodichogamy systems (Groh et al. 2025b, Groh et al. 2025a, Liu et al. 2025, Zhao et al. 2025) and many plant sex determining regions (e.g. Massonnet et al. 2020, reviewed in Renner and Müller 2021).

To test whether A1 and A2 are differentially regulated in additional species, we generated additional time courses of floral gene expression from Persea subg. Eriodaphne and Machilus thunbergii (Fig. 4). Overall, these data support that SDMYB expression peaks in correspondence with anthesis, and that the SDMYB A1 and A2 alleles have distinct diel expression patterns. In Persea subg. Eriodaphne, data from two A-type Persea caerulea (one homozygote and one heterozygote), a B-type of P. caerulea and a B-type of P. palustris support lower overall expression of the A1 SDMYB allele as seen in avocado. Our limited temporal sampling lacks power to detect a phase difference, but the data are nonetheless consistent with this possibility. We sampled flowers of an A-type Machilus thunbergii at higher temporal density and recovered a phase shift similar to that seen in avocado - a 2.57 hr phase delay in the A1 allele (P = 6.49 ⇥ 10*^—^*^4^). Here, the A1 allele does not exhibit lower average expression, but it does show significantly larger amplitude (P = 5.97 ⇥ 10*^—^*^3^). Taken together, these results indicate that conserved cis-acting regulatory differences between the SD-locus haplotypes have similar effects in avocado and its wild relatives of replicates.

**Figure 4:**
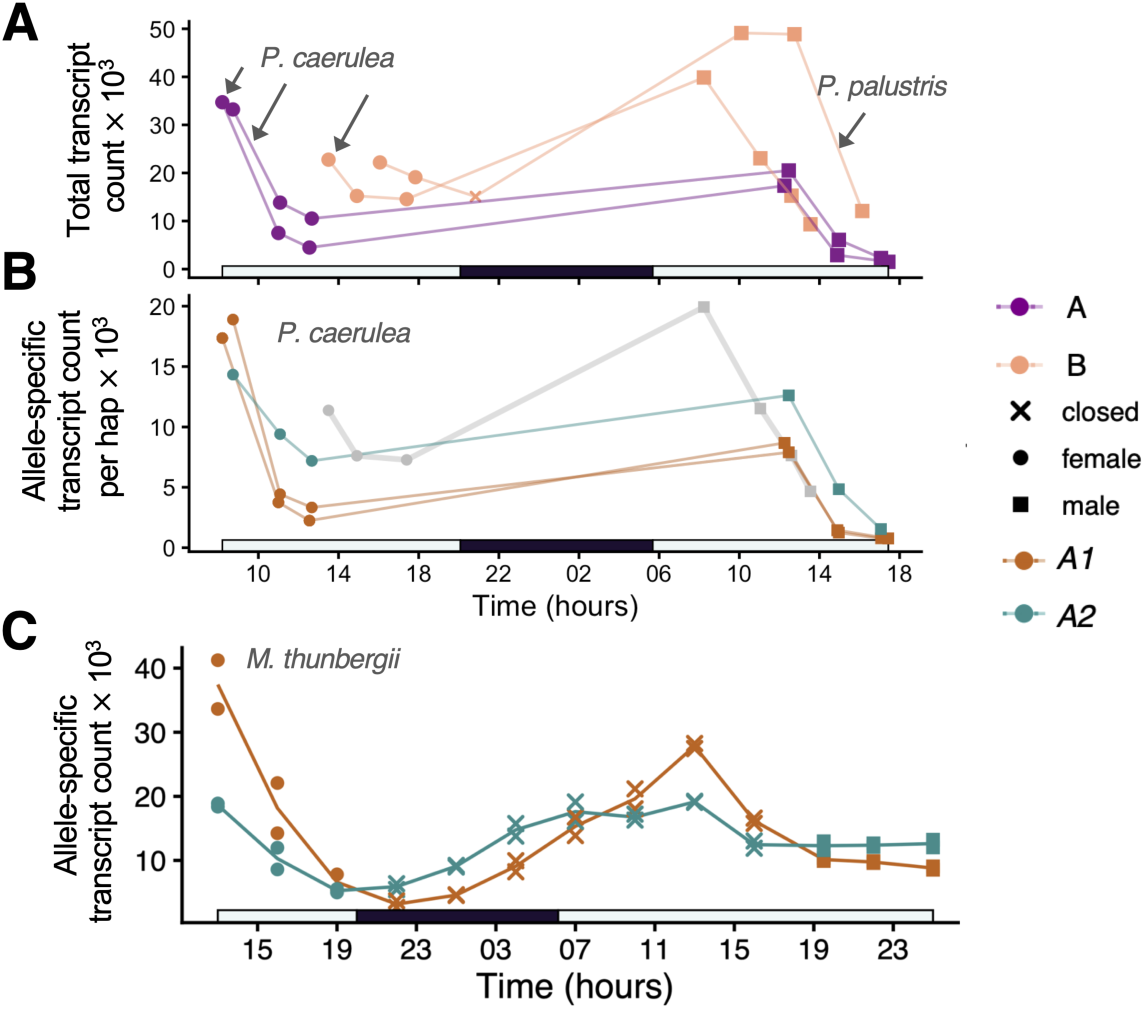
Conserved allele-specific regulation of *SDMYB* across Perseeae. **A)** Time series of *SDMYB* transcription for three individuals of *P. caerulea* and one individual of *P. palustris*. The A-type *P. caerulea* individual showing higher average expression of *SDMYB* is an *A1/A2* heterozygote while the other is a *A1/A1* homozygote. **B)** Time series of allele-specific *SDMYB* expression for the *P. caerulea* individuals from (A). Transcript counts are normalized to haplotype dosage to facilitate comparison of *A1* expression between the homozygote and heterozygote. Gray line shows one half the expression of the *A2* allele seen in the B-type individual from (A). **C)** Time series of *SDMYB* allele-specific expression for a single *M. thunbergii* A-type heterozygote. Each point represents a separate flower and lines show average.

## Discussion

We have shown that synchronized dichogamy in avocado its wild relatives is linked to diel expression rhythms of two ancient alleles of the floral transcription factor SDMYB. Our findings provide a genetic explanation for a century-old mystery of avocado pollination, anticipated by Stout (1927) who predicted that “such decided and specific differences in action can only be due to differences in the inherent constitution of the two groups of varietes.” Several aspects of the regulatory mechanism nonetheless remain to be solved.

First, it seems likely that the conserved non-coding polymorphisms within the SD-locus haplotypes mediate differential responses to an upstream transcriptional regulator with rhythmic activity. How do different rhythms of SDMYB alleles result from differential responses to transcriptional regulators? As the avocado flowering rhythm is at least in part circadian (Sedgley 1985), and SG19 MYBs are transcriptionally regulated by clock-regulated jasmonate signaling components (Mandaokar et al. 2006; Cheng et al. 2009; Reeves et al. 2012; Schubert et al. 2019; Thines et al. 2019), similar signaling components may be involved here. Separately, morph-specific disruptions to the avocado flowering rhythm by light and temperature (Sedgley 1977, 1985) suggest the diel expression difference of SDMYB alleles may be contingent on environmental cues. Consistent with this idea, some SG19 MYBs have been found to be light- and heat-sensitive (Shin et al. 2002; Wu et al. 2021). The observed diel expression difference between alleles could also result in part from transcriptional regulation of one allele by the other, either directly or indirectly.

Second, while the delayed rhythm of the A1 allele can explain the delay of 2nd anthesis of A-types, it is not sufficient to explain the earlier 1st anthesis of A-types. Thus, we speculate that the offset of the female and male flowering phases in opposing directions between flowering types derives from a combination of the differential SDMYB rhythmicity following 1st anthesis, as shown here, together with different allele affects on development in response to an additional time-varying factor. Because our developmental time course does not include flower buds prior to their first anthesis, we cannot exclude the possibility that the observed phase shift is specific to the period following the onset of 1st anthesis.

Expression differences alone are unlikely to be the sole basis of synchronized dichogamy in these species. The trans-species protein-coding differences between the haplotypes, in otherwise highly conserved domains, suggest that the SDMYB alleles may have partially diverged in function. These amino acid differences might modulate a differential response to fluctuating concentrations of hormone signaling components or transcription-activation binding partners (Song et al. 2011; Qi et al. 2015; Huang et al. 2017, 2020). Given that the expression of A2 is not markedly changed in the heterozygote background, particularly across second anthesis (Fig. 2D), differential protein-protein interactions could potentially form the mechanistic basis of dominance of the A1 allele. One candidate set of binding partners are the JAZ proteins, jasmonate-responsive negative regulators that directly interact with SG19 MYBs to regulate flower maturation in Arabidopsis (Song et al. 2011). Interestingly, some avocado homologs of the Arabidopsis JAZ proteins exhibit different expression rhythms between A- and B-types, implying these are regulated by SDMYB activity (Fig. S7A). Because each flower must open and close twice, some such negative feedback loop could be a core component to the regulation of this mating system and biphasic flowering rhythms more generally.

Disassortative mating generated by heterodichogamy is hypothesized to drive strong negative frequency-dependent selection pushing populations towards a 1:1 morph ratio (Gleeson 1982). The few natural samples of Perseeae examined to date suggest the two morphs occur in roughly equal proportions (Skutch 1945; Watanabe et al. 2016; Rogers 2024) (see also Fig. 3D), consistent with this prediction. Our identification of diagnostic trans-specific SNPs at the SD-locus now enable the estimation of morph frequencies and the strength of disassortative mating in a broader phylogenetic context. As synchronized dichogamy appears highly environmentally sensitive in avocado (Sedgley 1977, 1985; Pattemore et al. 2018), it will be interesting to measure variability in the strength of disassortative mating across the Perseeae tribe, and explore whether buffering mechanisms have evolved in species that naturally experience a high degree of environmental variation.

Our comparative analyses of the Perseeae SD-locus further establish heterodichogamy as a powerful agent of long-term balancing selection. Clear examples of long-term balancing selection remain relatively rare in plants, and are most often associated with mating type loci and and immune genes (Bechsgaard et al. 2006; Koenig et al. 2019). The SD-locus alleles likely diverged in the ancestor of Perseeae likely over 44 million years ago, and appear to be common among the hundreds of species in Perseeae (Fig. 3C). The age of this system is comparable to the ages of the several genetic systems for heterodichogamy in Juglandaceae (independently balanced for >40 Myr,Groh et al. 2025b, Groh et al. 2025a). It is noteworthy that these estimates are comparable with the oldest reported angiosperm sex chromosomes (e.g. Cherif et al. 2013; Carey et al. 2025, reviewed in Renner and Müller 2021), despite dioecy having evolved hundreds or thousands of times, whereas heterodichogamy is considerably more rare. Dioecy can be susceptible to rapid reversion to hermaphroditism under mate limitation (Cossard et al. 2021). By contrast, the hermaphroditic condition of heterodichogamous plants may afford greater reproductive assurance and long-term stability of the causal polymorphisms. On the other hand, one way heterodichogamy might be lost is through disruptive selection on sex allocation and the evolution of dioecy (Pendleton et al. 2000; Pannell and Verdú 2006). The presence of unisexual flowers in Alseodaphnopsis (Van der Werff 2019), which our analysis shows is nested within Perseeae (Fig. 3A), suggests a promising system for investigating this evolutionary pathway to dioecy.

Although a highly similar form of synchronized dichogamy is found in tribe Cinnamomeae (Hathurusinghe et al. 2023), our analyses indicate the mating system in Cinnamomeae is not controlled by the same set of alleles. In conjunction with the recent identification of the genetic control of flexistly in tropical gingers (Zhao et al. 2025), our results indicate that there are multiple developmental and gene regulatory avenues by which daily forms of heterodichogamy can evolve. As protogynous dichogamy is ubiquitous in magnoliids with cosexual flowers, and 2-day flowering rhythms linked to the day/night cycle are common (Endress 2010), magnoliids may be predisposed to evolve daily forms of heterodichogamy, known in at least four families of this group (Teichert et al. 2011; Endress 2020). We suggest that unlike seasonal forms of heterodichogamy, evolution of heterodichogamy over daily timescales is more likely to involve genes with late-acting roles in flower maturation. As SDMYB homologs are key regulators of diverse aspects of floral maturation (Reeves et al. 2012; Chopy et al. 2023), we speculate that convergent evolution of daily heterodichogamy may often recruit genes closely interacting with SDMYB, or perhaps even homologs of SDMYB itself. Interestingly, the avocado ortholog of SMPED1, a gene controlling flexistyly in Alpina mutica (Zingiberaceae), shows differential regulation between flowering types at second anthesis (Fig. S7B), consistent with this idea and its suggested role in anther dehiscence (Zhao et al. 2025). As these pathways become increasingly well characterized, this knowledge will enable us to further define the genetic factors that promote or constrain the evolution of diverse mating strategies, and allow for genetic manipulation of key flowering traits in important agricultural crops.

Finally, the identification of the genetic basis underlying A- and B-type flowering behavior has direct implications for avocado breeding. Due to the protracted juvenile phase of avocado (5-12 years, Lavi et al. 1992), breeding programs face long delays evaluating segregating populations and selecting parental lines. The genetic markers identified here will enable rapid screening for flowering type in seedlings, substantially reducing time, labor, and costs associated with field evaluation. Our findings further suggest SDMYB as a key target for directed breeding efforts, for instance to develop new pollenizers of ‘Hass’, the most widely planted and consumed avocado cultivar worldwide. Further genetic dissection of the avocado flowering mechanism will enhance strategies for breeding and orchard design, improving yields and sustainability of a globally significant crop.

## Acknowledgements

We are grateful to Michael Wenzel and other staff at Sonoma Botanical Garden, Holly Forbes and Sophia Warsh of the University of California Botanical Garden, Berkeley, CA, Ashraf El-Kereamy and Chris Martinez at UC Lindcove and South Coast Research and Extension Centers in Exeter and Irvine, CA, and Ross Brennan at UC Quail Ridge Reserve for facilitating data and tissue collection. Rebecca Gaut sampled and generated sequencing libraries for the avocado mapping panel. We thank Stacey Harmer, Veronica Thompson, Ben Blackman, Noah Whiteman, Jim Doyle, and members of the Coop lab for helpful discussions, and Michael Turelli for comments on an earlier draft of the manuscript. We acknowledge the DOE Joint Genome Institute for making public the Liriodendron tulipifera genome. Funding was provided by the National Science Foundation (NSF GRFP 1650042 awarded to J.S.G.), the National Institutes of Health (NIH R35 GM136290 awarded to G.C.), the UC Natural Reserve Maurer-Timm Endowment (awarded to J.S.G.), and NIFA AFRI (Grant no. 2021-67013-34237 awarded to M.L.A., D.S., B.G.). The authors declare no competing interests.

## Author contributions

J.S.G. and M.F.S.S. phenotyped plants; J.S.G. collected tissue samples with assistance from M.F.S.S., E.F., B.G., and G.C.; G.A. performed DNA and RNA extractions; J.S.G. and E.A.D. processed sequence data; J.S.G. performed data analysis and visualized results; E.F., E.S., D.S., B.G., M.L.A., and G.C. contributed data and resources; B.G., M.L.A., and G.C. provided supervision; J.S.G. wrote the manuscript with contributions from M.F.S.S., B.G. and G.C. All authors reviewed the manuscript.

## Data Availability

Genome assemblies of Persea americana ‘Puebla’, Machilus thunbergii, and Persea caerulea are available at NCBI through BioProjects PRJNA1392525-PRJNA1392526, PRJNA1392542-PRJNA1392543, and PRJNA1392566-PRJNA1392567, respectively. Supplementary Videos are available at https://doi.org/10.5281/zenodo.18021169. Raw assembly and resequencing reads will be made publicly available through NCBI.

## Materials and Methods

### Phenotyping

An open-pollinated orchard planted with ‘Gem®’ (patent no. 3-29-5, A-type heterozygote) and ‘Luna UCR®’ (patent no. BL516, B-type recessive homozygote) was established at University of California, Riverside prior to 1999, isolated a distance of 1/4 mile from other avocado orchards to favor crossing between these two varieties. Seeds from both maternal varieties were planted in an experimental plot over the course of several years from 2019-2022.

Phenotyping of the avocado mapping population was carried out at the UC Riverside Agricultural Experiment Station over four consecutive days during the first week of April 2024, coinciding with peak flowering in Riverside, CA. All individuals in bloom (n = 578) were visited twice per day, once in the morning (between 9:00 AM and 12:00 PM) and again in the afternoon (between 2:00 PM and 5:00 PM). Each visit targeted the detection of flowers at anthesis to determine whether the tree expressed female or male function during each time period. Plants were visited in reverse order on consecutive days. Due to suboptimal weather on the first two days, which may cause delayed flower opening (Pattemore et al. 2018), phenotyping was extended to a total of eight observation periods. This strategy allowed us to confirm consistent temporal sex expression patterns and assign each individual unambiguously to the A or B flowering type. Observations were made directly on the trees without the use of cameras, and phenotyping was timed to avoid confounding effects of early morning cold-induced floral delay or incomplete anthesis.

We examined flowering phenology of a natural population of California bay laurel Umbellularia californica at UC Quail Ridge Reserve in February 2025. Preliminary observations over the span of a day indicated that flowers of this species were not opening and closing within the span of a day. We then tagged up to 10 individual flowers on 17 individuals using jeweler tags and recorded the status of flowers over 5 separate visits in February. Flowers were considered in the female phase if the petals were at least partially open and the stigma appeared mature. Stigmas that appeared moist and whitish/translucent in color were judged as mature, and when they turned brown in color they were scored as finished. The male phase was considered as the period were at least one anther had dehisced, or all anthers had recently dehisced and pollen was still present.

### Association mapping

Sequencing reads of 712 avocado individuals were aligned to the Persea americana ‘Hass’ alternate assembly (GCA 031208655.1) (Nath et al. 2022) which we identified as containing the A1 haplotype in the initial stages of our study. This included the two parents and 710 putative progeny. After variant calling, genotypes at sites with GQ below 30 were set as missing. At this stage, 224 samples were filtered out because they had overly low coverage or a high proportion of variant sites that were not found in the two parents. We then performed imputation from the parental genomes using AlphaImpute2 (Whalen and Hickey 2020). Among the 486 imputed offspring, 374 were phenotyped for flowering type and were used in a Genome Wide Association Study (GWAS). GWAS was performed in gemma 0.98.5 (Zhou and Stephens 2012), removing sites with more than 5% missing data or less than 20% minor allele frequency in the sample. We calculated read depth from the alignment files across the GWAS association peak for all phenotyped individuals (including those with unexpected parentage that were phenotyped, N=504) using samtools (Danecek et al. 2021) and averaged read depth values in 1kb intervals. For each sample we then used the average depth across chromosome 9 to normalize windowed depth values.

In parallel, we re-examined published sequencing data from a sample of 23 avocado varieties and performed a GWAS using available phenotype information Solares et al. (2023). This analysis identified the same locus with highly similar patterns of segregating read depth, confirming that the association identified in the mapping population was not specific to the mapping population family (Fig. S1).

### Genome assemblies

We performed de novo genome assemblies for Persea americana ‘Puebla’ sampled from UC South Coast Research and Extension Center, Machilus thunbergii sampled from Sonoma Botanical Garden, Glen Ellen, CA (accession number 1987.421A), Persea caerulea sampled from plot 43, row 7, tree 1 at UC South Coast Research and Extension Center, and Persea podadenia from University of California Botanical Garden, Berkeley CA (accession number 97.0505.363). High molecular weight DNA extraction, library preparation, and of these four individuals across 2 PacBio Revio cells was performed by MedGenome, Inc., Foster City, CA. We used hifiasm 0.20 (Cheng et al. 2022) with default parameters to assemble CCS reads for each sample. For P. americana, we used ragtag (Alonge et al. 2022) with default parameters to scaffold the assembly into chromosomes using the avocado assembly of (Yang et al. 2024) as a reference. Measures of assembly contiguity are given in Table S3.

### Species phylogeny inference and time calibration

Relationships within the ca. 400 species of Perseeae remain unresolved, and previous molecular phylogenetic studies have typically relied on single genes (Rohwer et al. 2009; Li et al. 2011; Xiao et al. 2022; Chanderbali et al. 2024). Chen et al. (2020b) constructed a phylogeny of Lauraceae using 275 orthologs, but with limited taxon sampling from Perseeae. Furthermore, divergence time estimates for Perseeae have rested on either biogeographical assumptions (Chanderbali et al. 2001), or calibration using a single leaf fossil with uncertain taxonomic placement (Li et al. 2009, 2011), which may be problematic given the extent of homoplasy in Lauraceae leaf characters (Rubalcava-Knoth and Cevallos Ferriz 2025). To improve phylogenetic resolution in this group, we inferred a phylogeny for 32 Lauraceae taxa (Table S4) using 7,842 single-copy orthologs, and grafting this into a magnoliid backbone tree, used an alignment of 93 gene trees together with a set of fossil calibrations (Table S5) to time calibrate the tree (Table S6).

First, we identified a set of 7,862 single copy orthologs using fifteen genome assemblies spanning the core Lauraceae (tribes Perseeae, Laureae, and Cinnamomeae) (using OrthoFinder 2.5.5 (Emms and Kelly 2019) with option -d to run on DNA sequences). The input transcriptomes consisted of eight de novo genome annotations and liftoff annotations (Shumate and Salzberg 2021) from the avocado genome for unannotated assemblies. After initial ortholog detection, we then used shotgun resequencing data aligned to the P. americana genome to generate consensus sequences over these orthologs for an additional sixteen species. To do this, we first generated an all-sites vcf, then used bcftools consensus (Danecek et al. 2021), with parameters -e ‘FMT/GT==“./.”’ -a “N” -M “N” –mark-del “N” for each exonic region of each single copy ortholog, reverse complementing the result for sequences encoded on the reverse strand. Importantly, by treating sites with missing data as ‘N’ rather than assuming the reference allele (avocado), this approach avoids reference bias from impacting the phylogenetic placement of these samples. We converted any sites called as heterozygous within samples to missing data to avoid any impact of genotype priors and sequencing error in the variant calling step. Doing so, we recovered sequence from all species for 7,842 single copy orthologs, which were then used for phylogenetic inference.

Sequences for each orthogroup were aligned using muscle (Edgar 2004) and subsequently trimmed using trimal using the heuristic algorithm optimized for maximum likelihood phylogeny inference (Capella-Gutíerrez et al. 2009). We then inferred a gene tree for each orthogroup using maximum likelihood in iqtree with a GTR+I+G4 substitution model and 1000 ultrafast bootstrap replicates (Nguyen et al. 2015). Gene tree nodes with less than 10% bootstrap support were collapsed to polytomies (Zhang et al. 2018) using newick tools https://github.com/xflouris/newick-tools, and then we ran ASTRAL IV (Zhang et al. 2018) on the resulting gene trees to infer a species tree under a model that accounts for incomplete lineage sorting. In parallel, we used a concatenation+ maximum likelihood approach in IQTree using a GTR+I+R4 model with 1000 ultrafast bootstrap replicates. Notably, we recovered identical topologies, with all nodes showing LPP support values of 1 and 100% bootstrap support from these two methods, respectively. We rooted the core-Lauraceae trees at the edge separating Perseeae from Laureae-Cinnamomeae, as the relationships among tribes is well-established, and this was also confirmed in our magnoliid-wide analysis described below.

In parallel we inferred a backbone tree for magnoliids, in order to calibrate our core-Lauraceae tree using established calibration points across magnoliids. First, we used fourteen genome assemblies with independent annotations of protein-coding genes to define a set of 105 single copy orthologs across magnoliids, using OrthoFinder 2.5.5 (Emms and Kelly 2019) with coding sequences as input. These orthologs were roughly evenly distributed across all twelve autosomes of avocado. For eleven other genomes for which we could not access annotations, we used liftoff (Shumate and Salzberg 2021) to extract coding sequences of these orthologs, using a reference genome from within the same phylogenetic order and setting parameters ‘-mm2 options=“-r 2k -z 5000 –end-bonus 5” -d 5 -flank 0.2 -polish’ to improve alignments between distantly related species, resulting in sequences from at least 95 single copy orthologs for all taxa in our data set. This taxonomic sample spans all four orders within magnoliids and nine families. As described above, we inferred gene trees from trimmed alignments of each ortholog, and inferred species phylogenies using both ASTRAL and concatenation approaches. Importantly, the relationships among broader groups were in perfect agreement with maximal support using both methods, with the exception of relationships within core-Lauraceae, which we confidently resolved in the previous analysis. The tree was rooted along the edge separating the Canellales-Piperales clade from the Magnoliales-Laurales clade following Helmstetter et al. (2025).

We time calibrated the tree using the likelihood approximation method of MCMCTree (Yang and Rannala 2006). MCMCTree infers node ages using a fixed topology, which is appropriate as both our ASTRAL and concatenation+maximum likelihood analyses gave identical topologies. We specified priors on internal nodes with soft bounds to account for the possibility of fossil misplacement on the phylogeny. Due to the difficulty of unambiguously assigning fossils within Perseeae, we rely on a larger set of fossil calibrations previously established for magnoliids (Massoni et al. 2015a), together with additional calibrations for Lauraceae, which are detailed along with justification in Table S5. Briefly, our calibrations incorporate the wide uncertainty in the crown age of magnoliids (Sauquet et al. 2022; Zuntini et al. 2024), include established lower bounds for deeper nodes within magnoliids (Massoni et al. 2015b), and set priors on ages of nodes within Lauraceae that are consistent with patterns of diversification of this group in the late Cretaceous and Cenozoic (Taylor 1988; Doyle and Endress 2010; Friis et al. 2011; Huang et al. 2016) and that are consistent with estimates from previous studies (Li et al. 2011; Huang et al. 2016; Xiao et al. 2022).

For nodes without specified priors, the priors are defined based on the birth-death model with the default parameters set by the program, which gives a uniform distribution of node ages for uncalibrated nodes, subject to constraints of the use-calibrated nodes. Ninety-three of the 105 single-copy orthologs from our magnoliid-wide orthology search were also contained in the set of single-copy orthologs defined for core Lauraceae. We therefore concatenated the alignments of these genes for the molecular data input to MCMCTree. We ran MCMCTree with a correlated rates relaxed clock model and the HKY85+G5 substitution model. We specified a diffuse prior for substitution rates with the gamma rate parameter ff = 2 as recommended by the program manual. To set the mean of the prior on substitution rates, we used the mean too to tip height from the maximum likelihood tree of magnoliids together with the mean of the prior on the root age to derive an estimate of substitutions per site per 100 million years. We first ran MCMC algorithm by sampling from the prior to verify that our prior specification was sensible. We ran two separate chains of the MCMC algorithm with substitution data for a sample size of 50,000, sampling every 10 iterations after a burn-in period of 5,000 iterations. Results were checked for convergence across multiple chains.

### Gene annotation and structure of the SD-locus

Using the available gene annotation for ‘Hass’ (Nath et al. 2022), we detected conserved gene synteny between the two haplotypes using reciprocal BLAST searches (Fig. 3E). We also manually screened the region to examine evidence for the expression of other unannotated genes using our floral expression data, but none were detected. We did note that the published annotation for the gene MRB53 031467 appeared to be missing two exons which we included in alignment of reads to transcriptomes for later analyses. We used InterPro scan and BLAST searches to investigate gene identity. We extracted regions homologous to the SD-locus identified in avocado and generated dotplot alignments using the online version of megablast through NCBI (Fig. S4). Coordinates of syntenic genes were identified through command line blastn. We annotated each assembly using EDTA v2.2.2 (Ou et al. 2019) and plotted the locations of repeat annotations along the coordinates of these alignments.

### Analysis of SDMYB homologs

We first used blastp to identify genes with close homology to SDMYB in other species including Arabidopsis. This identified SDMYB as a member of the SG19 subgroups of transcription factors (Fig. S2A). We produced an alignment of amino acid sequences using muscle (Edgar 2004) for a set of SG19 R2R3 Myb transcription factors with previous functional characterization in addition to close outgroups to the Perseeae haplotypes (Fig. S2A).

To infer relationships among Lauraceae sequences at the SD-locus, inferred a phylogeny from separately aligned regions spanning the region of trans-species polymorphism. This included (i) approximately 6 kb upstream of the start codon of SDMYB, (ii) the coding sequence of SDMYB, (iii), intron 2 (approximately 6 kb), and a ⇠5 kb region downstream of SDMYB distal to the structural variant. For the coding sequence of SDMYB, we performed a codon-aware alignment and truncated the alignment after the frame shift in the final exon. For the other three non-coding regions, we used blastn to identify coordinates of homologous regions from each assembly, extracted these using samtools (Danecek et al. 2021), reverse complementing as needed, then used muscle (Edgar 2004) to align these regions. Our genome assembly from an A-type P. caerulea individual revealed that it was a homozygote for the A1 haplotype, thus we were lacking an A2 sequence for P. caerulea. Therefore, we supplemented our genome sequences by extracting exonic SDMYB sequence from a B-type individual of P. caerulea for which we had performed RNAseq. To do this, we aligned the transcripts to avocado genome using STAR (Dobin et al. 2013) and then used bcftools (Danecek et al. 2021) to extract a consensus sequence from over the exons separately for each sample. Coverage was very high over the exons for the samples used, and we first filtered variants for GQ 40, then combined these with invariant sites. Any sites with missing data were coded as ‘N’ rather than assuming the reference sequence. We also aligned to the dominant allele of avocado, whereas this B-type sample has two recessive alleles. Thus, reference bias does not impact the phylogenetic placement of these samples. We used a similar procedure to incorporate Cinnamomum verum into the phylogeny by extracting a consensus sequence from WGS alignments over the SDMYB coordinates in the avocado genome. We trimmed the alignments using trimal (Capella-Gutíerrez et al. 2009), and further curated each alignment by removing portions of the alignment where homology could not be confidently assigned. Following this, we fit an edge-proportional partition model in IQTree (Nguyen et al. 2015), specifying a MG+F1X4+G4 codon substitution model (Muse and Gaut 1994) for the coding sequence of SDMYB and the HKY+F+G4 nucleotide substitution model (Hasegawa et al. 1985) for the non-coding regions. We rooted the tree by specifying Sextonia rubra as the outgroup. We calibrated the maximum likelihood tree using penalized likelihood implemented in the chronos function in the R package ape (Paradis and Schliep 2019). We used the divergence of Perseeae from Cinnamomeae and Laureae as a fixed calibration point and assumed a strict clock model. We used the posterior mean estimate obtained from our species tree, as well as the 95% HPD lower and upper bounds to obtain confidence intervals for the date estimate of the SD-locus haplotypes.

To measure nucleotide divergence of core Lauraceae genomes against the genome of Persea americana ‘Hass’, we used AnchorWave (Song et al. 2022) with settings -R 1 -Q 1 to align against the ‘Hass’ alternate assembly (A1 haplotype), and used a custom R script to calculate nucleotide divergence in 1 kb windows in coordinates of the reference. For visualization, we removed windows with either fewer than 500 aligned base pairs in the reference and fit a LOESS smooth curve to the data with span=0.75, separately on either side of the large structural variant. We also filtered the values immediately adjacent to the structural variant which were unrealistically high and we considered to result from alignment error.

### Gene expression sampling design and analysis

We studied temporal patterns of gene expression in relation to floral anthesis and sex expression in Persea americana, Persea caerulea, and Machilus thunbergii, representing three divergent clades within Perseeae (Fig. 3A). In Persea americana, we sampled three biological replicates (three individuals that were not clones) at regular three hour intervals over a 21 hour period on April 11th, 2024 from the same plot at UC Lindcove Research and Extension Center in Visalia, CA. On the previous day, we marked numerous individual flowers on each tree that opened during first anthesis using jeweler tags, indicating their day of opening. Sampling began at 4:30 AM the following morning and continued until 1:30 AM the following day. Individual flowers were removed at the receptacle using tweezers, placed directly into a 1.5mL tube, and immediately flash frozen in liquid nitrogen, recording the exact time to the minute. Note all sequenced samples represent RNA extracted from single flowers. At each time point, trees were sampled in a consistent order, which alternated between A and B types. At each sampled time point, the total time to sample all 6 trees was approximately 10-15 minutes. Flowers that opened during first anthesis on the sampling day were also tagged such that they could be sampled after closing. Sampling stopped when flowers had closed after their second opening with the exception of two flowers of B-types. We also added an additional time point at the 2nd anthesis of B-types as our regular 3 hour intervals would have otherwise missed this developmental landmark. During the main sampling day and for to two nights previous, we used WingScapes BirdCam cameras to take photographs of inflorescences at regular intervals every 10 minutes. Photographs were subsequently stitched together to create time lapse videos of flowering (see Supplementary Videos).

Samples for the expression time series of Persea caerulea and P. palustris were sampled from Field 43 at UC South Coast Research and Extension Center in Irvine, CA on June 21, 2024. In this case, we sampled three time points during each of the two flowering phases in a single day. One of the A-type individuals sampled was the same individual used for genome assembly.

Samples for the expression time series of M. thunbergii were taken from a single individual with A-type flowering on May 2, 2025 at Sonoma Botanical Garden, Glen Ellen, CA. We used time-lapse photography to confirm the flowering type (Supplementary Video 4), and this individual was the same as used for our genome assembly.

RNA was extracted from whole flowers using the Qiagen RNAeasy Plant kit. Library prep and Illumina sequencing was performed by Signios Bio, Inc., Foster, CA (formerly MedGenome).

We quantified transcript counts using salmon (Patro et al. 2017). Raw RNAseq reads were first trimmed using skewer (Jiang et al. 2014), and then aligned to the transcriptome extracted from the ‘Hass’ genome annotation (Nath et al. 2022) downloaded from NCBI. To quantify both total and allele-specific expression of these three genes through time, we included both alleles found on the dominant and recessive haplotypes in the reference transcriptome. This approach allows RNAseq reads to align to either allele, avoiding the known confounding effects of reference bias on transcript quantification. To improve mapping quality for these three genes, we used the RNAseq alignments to manually edit the transcript sequences to include UTR sequence, and also in doing so identified that the ‘Hass’ annotation for MRB53 031467 was missing two exons, which we included in the reference for alignment.

The allele-specificity of this quantification approach depends on the density of allele-specific SNPs in the reference transcriptome copies. In total in avocado, we found 26 SNPs differing between the A1 and A2 haplotypes across the three exons of SDMYB, 9 SNPs across the 6 exons of MRB53 031467, and 3 SNPs across the 8 exons of MRB53 031468. Within the mature transcript sequence of SDMYB, the largest distance between two haplotype-specific SNPs is 148 bp, and each member of a read pair is 150 bp. Thus, the vast majority of reads will overlap at least one haplotype-specific SNP and we expect our quantification of allele-specific expression to be accurate. In genes MRB53 031467 and MRB53 031468, the longest portion of the transcripts containing no haplotype-specific SNPs are approximately 670 bp and 950 bp, representing approximately 49% and 54% of the total transcript lengths. The majority of reads are expected to overlap at least one SNP, but those that don’t could plausibly align with equal probability to either of the two alleles, making quantification of allele-specific expression conservative. Thus, as a complementary approach, we separately quantified allele-specific expression by aligning all reads to the same reference and directly measuring allelic depth at haplotype-specific SNPs, taking a depth weighted average of relative allele depth across SNPs. We found highly similar results using both approaches and report the results from the competitive mapping.

For P. caerulea, P. palustris and M. thunbergii we used liftoff (Shumate and Salzberg 2021) to port the ‘Hass’ annotation to these assemblies, then extracted the corresponding coding sequences for the reference transcriptome. To measure expression of the duplicate copies of MRB53 031467 present in the P. caerulea dominant haplotype we used splice-aware aligned RNAseq reads to manually extract the coding regions corresponding to both copies. Our P. caerulea assembly was from a dominant homozygote individual, so to obtain the coding sequence of SDMYB for the recessive haplotype, we used the short read RNAseq data from the B-type individual of P. caerulea. Here, we extracted the consensus sequence over the exons and randomly resolved any heterozygous present sites to one allele.

For comparison across samples, we obtained transcript counts normalized to library size using the normalized count model of DESeq2 (Love et al. 2014). For the three genes where we considered alleles separately, we summed the normalized counts across alleles to obtain a gene-level normalized transcript count.

We used Principal Components Analysis (PCA) to examine the major axes of gene expression variation across all samples. To remove any variance associated with individual-level effects, we used redundancy analysis in the R package vegan to first remove axes of variation associated with individual replicates by using a design matrix to specify individual of origin (this had little effect on the first two axes of unconstrained variation). PCA was run for all samples combined (Fig. 2A), and separately for only open flower samples, (Fig. 2B) in order to better resolve expression variance between male and female phases.

To test for a morph difference in overall expression levels of SDMYB, we fit a mixed effects model in the R package lmer, specifying sampling time point (as in Fig. 2) and individual as random effects. We used a similar procedure to test for a difference in the average expression of the two alleles in heterozygotes. We fit a cosinor regression model to the SDMYB alleles in avocado using the R package CircaCompare (Parsons et al. 2020) to estimate the phase shift between alleles and test for statistical significance.

### Detection of *SDMYB* polymorphism across species

To examine the segregation of the SD-locus haplotypes across Perseeae, we genotyped samples at diagnostic SDMYB exonic SNPs in public resequencing data from published studies, including low coverage data (⇠2x) (Rendón-Anaya et al. 2019; Chen et al. 2020b; Tian et al. 2021; Bai et al. 2022; Xiao et al. 2022; Xiaoxuan et al. 2022; Zhu et al. 2023; Solares et al. 2023). We first defined exonic, biallelic, trans-specific SNP sites within SDMYB. We identified 2 in exon 2 (one is the nonsynonymous substitution in the R2R3 domain) and 2 in exon 3. The SNPs in each exon are 47 and 15 bp apart, typically spanned by single reads, whereas these sets of SNPs are 6.5 kb apart and should not be spanned by correctly aligned read pairs in these data. We selected one of these SNPs per exon, and obtained allelic depth at these SNPs from alignment files. We calculated genotype likelihoods using combined allele counts at the two SNPs with a binomial probability model allowing for a 1 ⇥ 10*^—^*^3^ probability of sequencing error. Using an equal prior probability for each of the three genotypes, we then computed genotype posterior probabilities (Table S7).

If a sample showed zero or only one read covering the chosen SNP site in either exon, we used read counts from the neighboring ascertained diagnostic site within the same exon if the depth was higher at that site. The frame-shift mutation in the final exon is also conserved across Perseeae, and we noted that samples with high posterior probability of being heterozygote based on these SNPs also showed the frame shift polymorphism with sufficient depth (Fig. S5). For any individuals polymorphic for the complex frame-shift mutation, which we considered highly unlikely to result from sequencing error (mutation from one allele to the other requires at least 2 insertion/deletion mutations), we set the genotype posterior to be 1 for the heterozygote.

To visually demonstrate that the segregation of TSP SNPs in individuals in additional species of unknown phenotype was restricted to SDMYB, we visualized SNP sharing with Persea americana ‘Hass’ in exonic regions for a genomic window surrounding SDMYB with multiple genes on either side (Fig. S5).

