## Supplementary_Figures for "Balanced polymorphism in a floral transcription factor underlies an ancient rhythm of daily sex alternation in avocado"

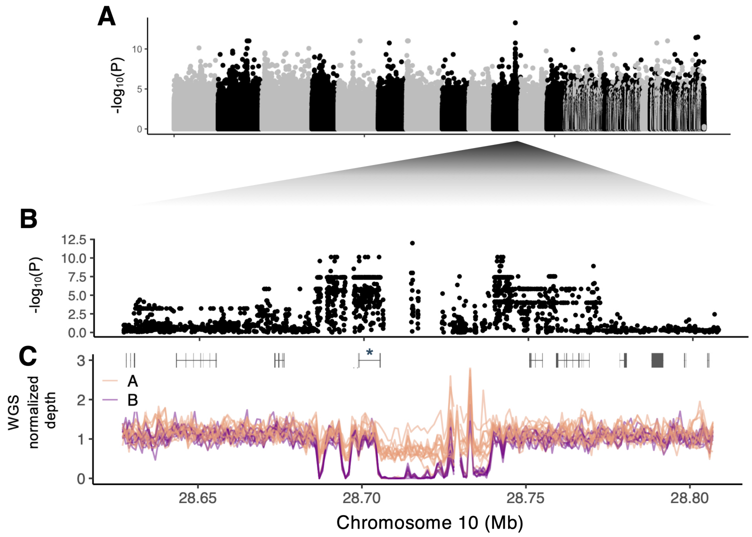


Figure S1: **A)** GWAS using published data (Solares *et al.* 2023; Rendón-Anaya *et al.* 2019) of 23 avocado varieties not in our mapping population against an assembly of 'Gwen' (Solares *et al.* 2023) **B)** Zoomed in view of the GWAS peak region in the 'Hass' alternate assembly (Nath *et al.* 2022) with gene models at bottom. **C)** Normalized read depth in 1 kb windows across this region for the same samples.

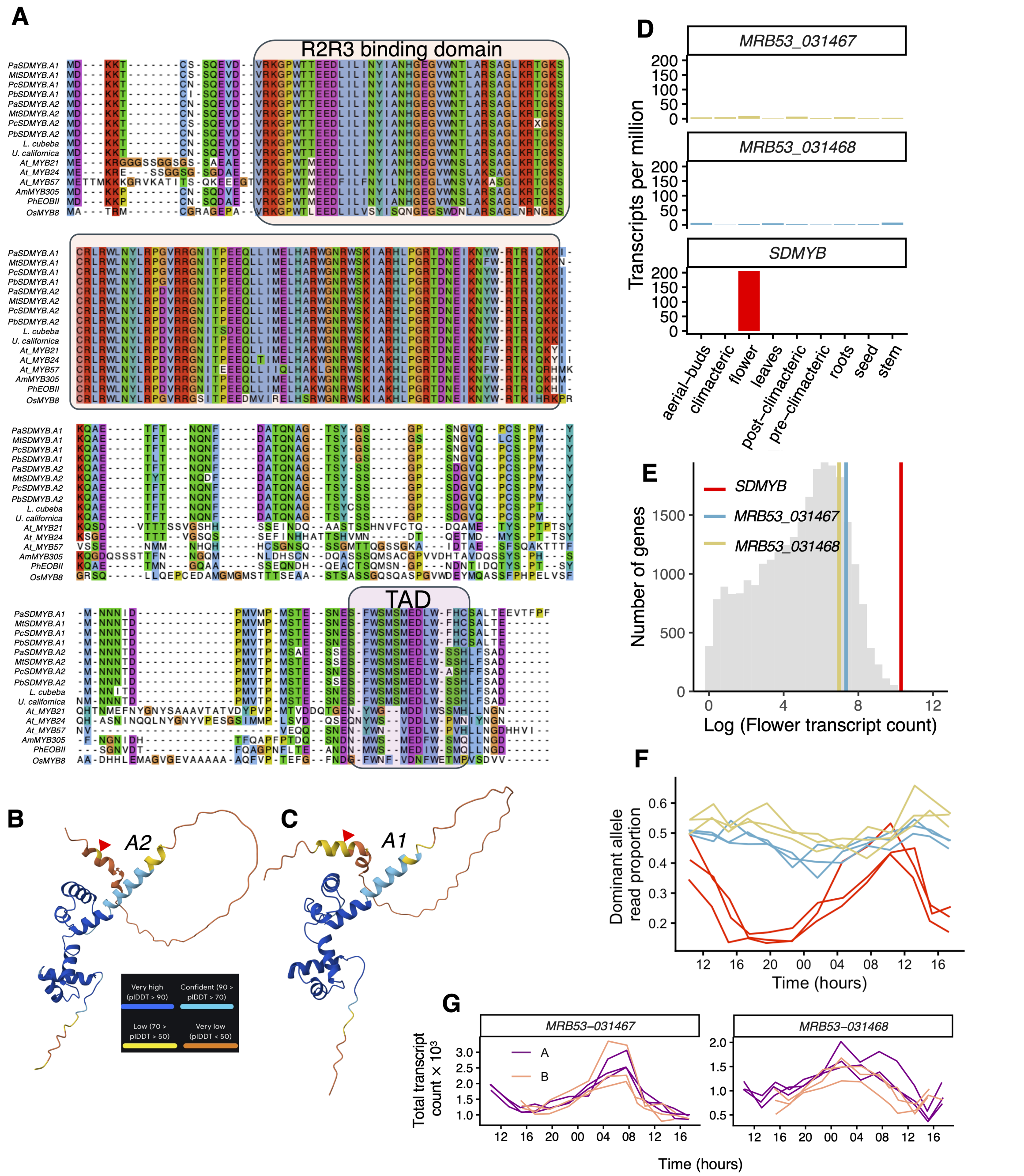


Figure S2: **A)** Alignment of SG19 R2R3 amino acid sequences. Species abbreviations in sequence names as follows: Pa=*Persea americana*, Mt=*Machilus thunbergii*, Pc=*Persea caerulea*, Pb=*Phoebe bournei*, followed by *Litsea cubeba* and *Umbellularia california*, followed by At=*Arabidopsis thaliana*, Am=*Antirrhinum majus*,Ph=*Petunia hybrida*, Os=*Oryza sativa*. **B,C)** AlphaFold predictions of protein structure for the *A1* and *A2* avocado alleles. The R2R3 DNA binding domain is highly structured (dark blue). The TAD is in an unstructured region of the protein but has a predicted alpha-helix structure. Location of frame-shift mutation indicated with red triangles. **D)** *SDMYB* shows flower-specific expression, unlike the other two genes near the association peak. Data are from several tissues from a single avocado individual (Ibarra-Laclette *et al.* 2015) used elsewhere for genome assembly (Rendón-Anaya *et al.* 2019), whose genotype we confirmed as *A2/A2* from published genomic sequence. **E)** Average expression level of all flower-expressed avocado genes across 73 samples (Table S2). The expression level of three genes at the SD-locus are highlighted with colored vertical lines. **F)** Proportion of transcripts derived from the dominant haplotype for three genes at the SD-locus association region. Colors are the same as in (E). **G)** Total transcripts by flowering type of the two additional genes besides *SDMYB* found at the SD-locus association peak.

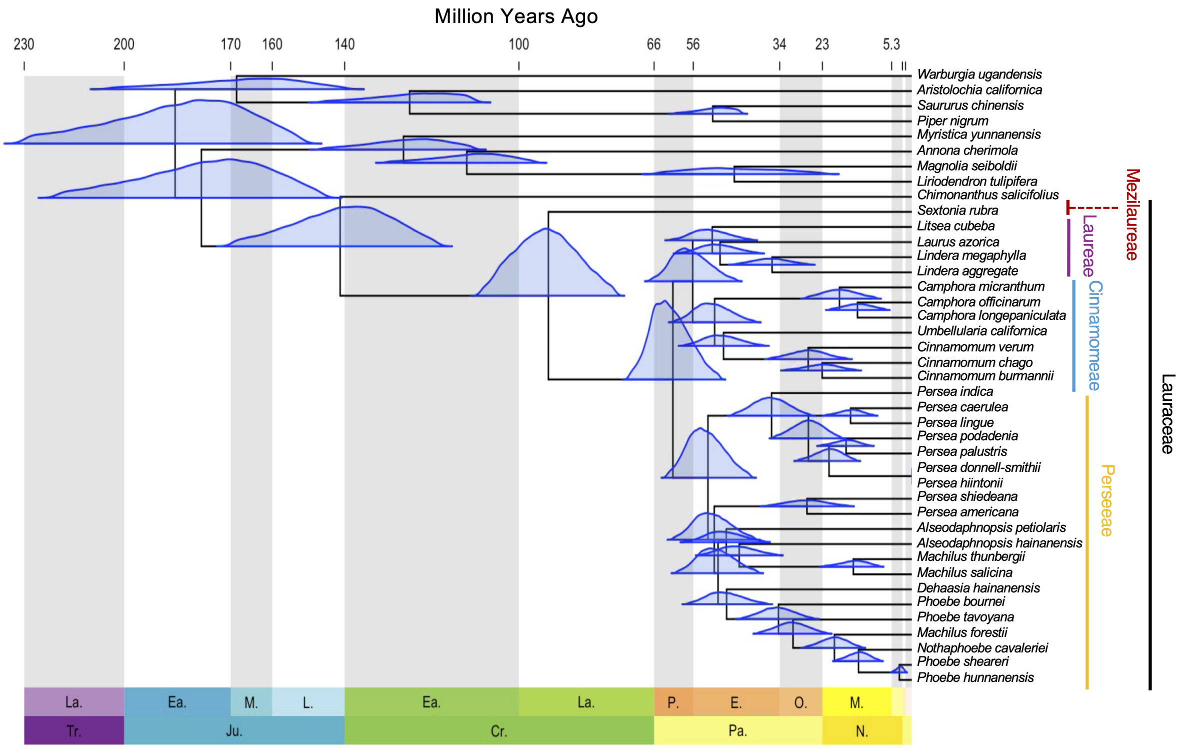


Figure S3: **A)** Fossil-calibrated phylogeny of a subset of Magnoliids. Taxon sampling is targeted to Lauraceae and the inclusion of at least one members of each magnoliid order, but is not meant to be exhaustive and reflects availability of public whole genome sequence data suitable for our analysis method. The blue distributions are posterior probability distributions from MCMCTree (see Methods, point estimates and credible intervals in Table S6). See Table S5 for fossil calibrations. Not all Lauraceae tribes are shown, see Li *et al.* (2025) for a review of broad level taxonomy in Lauraceae. All nodes in the tree shown had maximum support in both ASTRAL and concatenation + maximum likelihood methods.

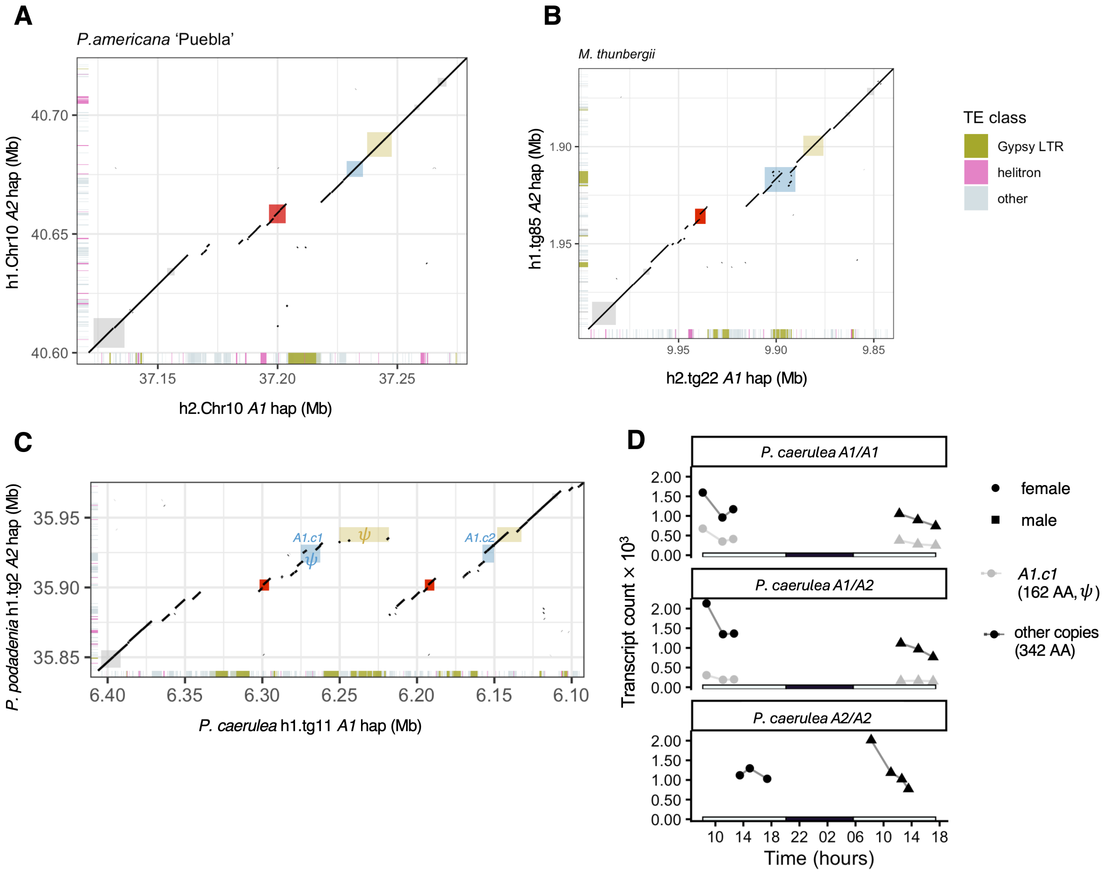


Figure S4: **A-C)** Dotplot alignments between alternative SD-locus haplotypes within *P. americana* (A), *M. thunbergii* (B), and *Persea* subg. *Eriodaphne* (C). Syntenic gene regions are highlighted in center with colored coxes, *SDMYB* in red. Transposable element annotations are shown at edges with two classes indicated in color. In (C), $\psi$ denotes predicted pseudogenes in the *A1* haplotype of *P. caerulea*. Two duplicate copies of *MRB53-031467* are labeled in blue, see also panel (D). **D)** Expression of two paralogs of *MRB53-031467*, the gene downstream to *SDMYB* copies within the dominant haplotype of *P. caerulea*. Panels represent different individuals of *P. caerulea*. We were unable to differentiate between the transcripts from the recessive haplotype copy of this gene and the 2nd copy found on the dominant haplotype, *A1.c2*, so transcript counts from these are pooled. However in the *A1/A1* homozygote (top) we can be sure this pool represents only the *A1.c2* copy. In this individual (top), *A1.c1* shows low expression and has a truncated amino acid sequence, appearing to be a pseudogene.

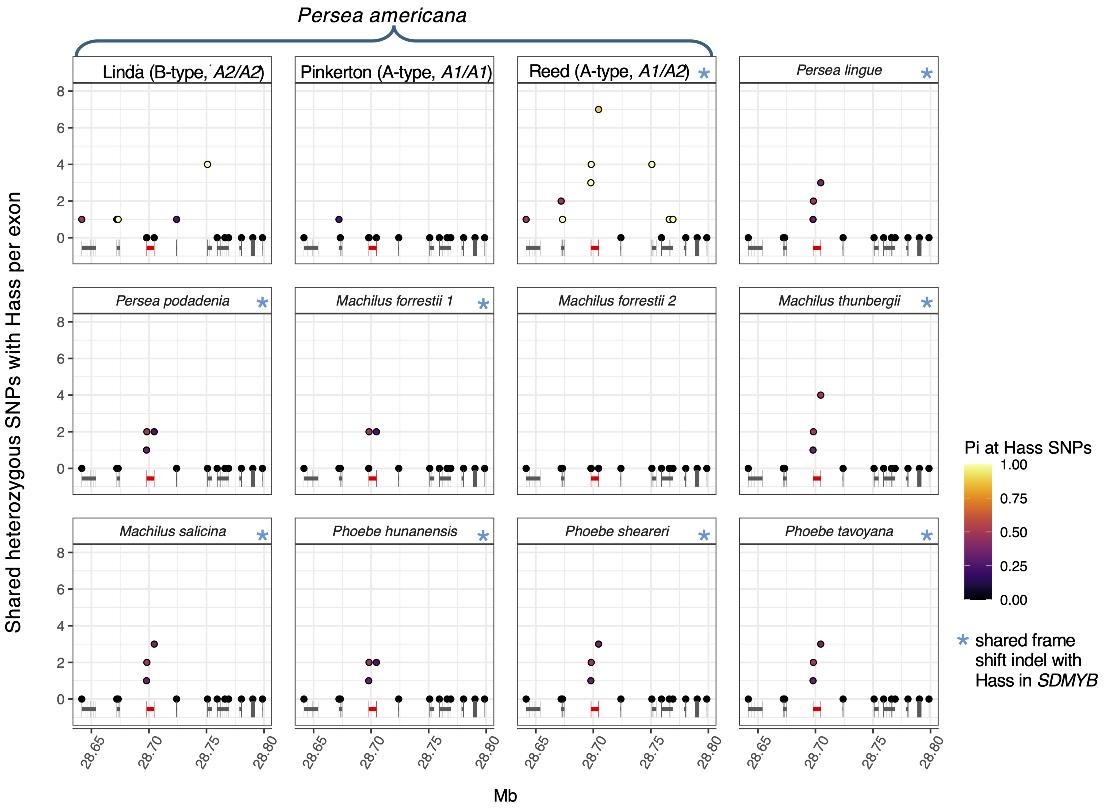


Figure S5: Shared heterozygous sites with *Persea americana* 'Hass' (A-type, *A1/A2*) in exonic sequence for a subset of the individuals examined (see Table S4 for full results). Each point represents a single exon, with the height of the point representing the number of sites that are shared SNP polymorphisms with 'Hass'. The color of each point represents the fraction of sites the individual is heterozygous at, out of the total number of heterozygous sites in 'Hass' for that exon. In addition, blue asterisk denotes the presence of a shared frame shift polymorphism in the last exon of *SDMYB*, which coincides with the presence of shared SNP polymorphisms.

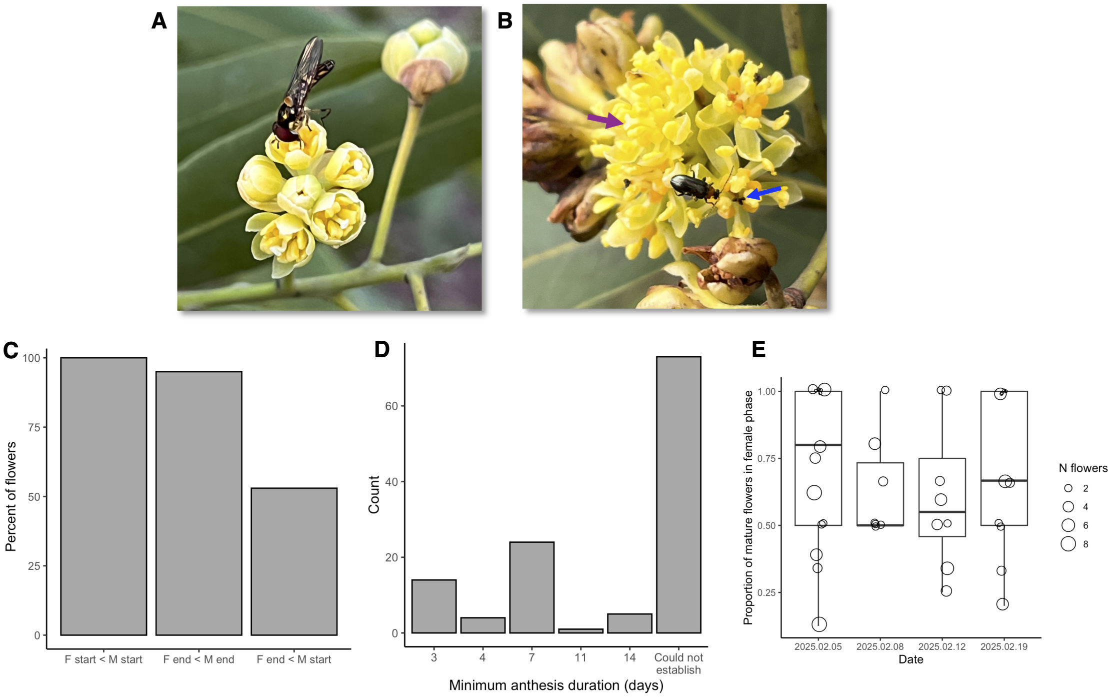


Figure S6: **A)** Flowers of *Umbellularia californica* in female phase. Stigmas appear moist and whitish/translucent, while anther valves have not dehisced. **B)** An example of overlap between female and male phase between flowers on the same plant. Purple arrow indicates a flower in female phase with a fresh stigma and undehisced anthers. Blue arrow indicates a flower with a dark whithered stigma and dehisced anthers. **C)** Summary of intrafloral dichogamy. All flowers are protogynous: the female phase always begins before the male phase and almost always ends before the male phase ends, with some degree of overlap based on visual assessment. F and M mark female and male respectively. **D)** A considerable proportion of flowers showed anthesis periods spanning more than several days. The x-axis gives the minimum number of days a tagged flower was inferred to be sexually mature over the multiple field site visits. This minimum couldn't be established if a flower went from unopened to completion of anthesis over two consecutive visits (the majority of cases), or if observations did not span the start and end of floral anthesis. **E)** Dichogamy is not synchronous between flowers on the same plant. Each dot shows a tree measured on a particular date, the size indicating the number of tagged, mature flowers open. See 'Phenotyping' section Methods for details.

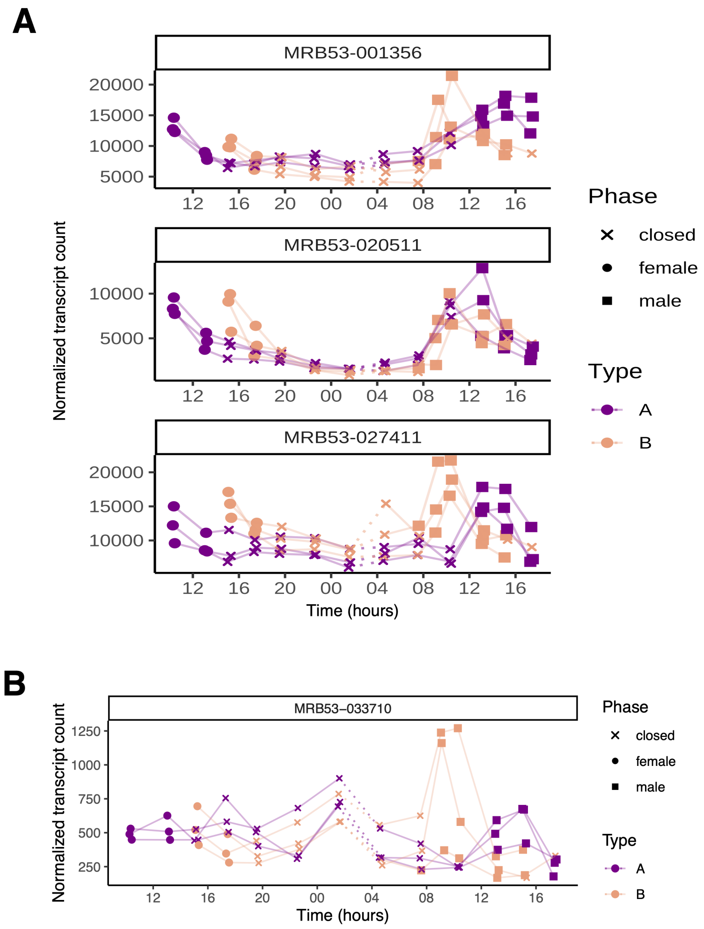


Figure S7: **A)** Several avocado homologs of the Arabidopsis *JAZ* genes show morph-specific expression patterns in avocado, suggesting that these are regulated by *SDMYB*. **B)** The avocado ortholog of *Alpinia mutica* Am*SMPED1* shows morph-specific expression, particularly at second anthesis.
